# A simple method to dramatically increase *C. elegans* germline microinjection efficiency

**DOI:** 10.1101/2023.03.23.533855

**Authors:** Theresa V. Gibney, Michelle Favichia, Laila Latifi, Taylor N. Medwig-Kinney, David Q. Matus, Daniel C. McIntyre, Angelo B. Arrigo, Kendall R. Branham, Louis T. Bubrig, Abbas Ghaddar, Juliana A. Jiranek, Kendra E. Liu, Charles G. Marcucci, Robert J. Porter, Ariel M. Pani

**Affiliations:** Department of Biology, University of Virginia, Charlottesville, VA, USA; Department of Biochemistry & Cell Biology, Stony Brook University, Stony Brook, NY, USA; David Matus is a paid consultant of Arcadia Science; Neuroscience Graduate Program, University of Virginia, Charlottesville, VA, USA; Department of Cell Biology, University of Virginia, Charlottesville, VA, USA

**Author notes:** Authors for correspondence Theresa V. Gibney, Ariel M. Pani. Department of Biology, University of North Carolina at Chapel Hill, Chapel Hill, NC, USA.

**Keywords:** *C. elegans*, microinjection, genome engineering, gene editing, CRISPR, transgenesis

## Abstract

Genome manipulation methods in *C. elegans* require microinjecting DNA or ribonucleoprotein complexes into the microscopic core of the gonadal syncytium. These microinjections are technically demanding and represent a key bottleneck for all genome engineering and transgenic approaches in *C. elegans*. While there have been steady improvements in the ease and efficiency of genetic methods for *C. elegans* genome manipulation, there have not been comparable advances in the physical process of microinjection. Here, we report a simple and inexpensive method for handling worms using a paintbrush during the injection process that nearly tripled average microinjection rates compared to traditional worm handling methods. We found that the paintbrush increased injection throughput by substantially increasing both injection speeds and post-injection survival rates. In addition to dramatically and universally increasing injection efficiency for experienced personnel, the paintbrush method also significantly improved the abilities of novice investigators to perform key steps in the microinjection process. We expect that this method will benefit the *C. elegans* community by increasing the speed at which new strains can be generated and will also make microinjection-based approaches less challenging and more accessible to personnel and labs without extensive experience.

## Introduction

A thorough understanding of many biological processes will depend on the ability to visualize cellular behaviors, subcellular structures, and protein dynamics in living systems. Researchers have available a wide range of genome manipulation methods to introduce transgenes and make insertions at endogenous loci in many traditional and emerging model systems. Depending on the experimental goals, these approaches can be used to both visualize native cell and protein dynamics and to introduce functional alterations to derive mechanistic insights. The nematode worm *Caenorhabditis elegans* is a model organism that is commonly used across a range of biological disciplines for its experimental tractability. *C. elegans* has a particularly well-developed repertoire of methods for transgenesis and genome engineering (Frokjaer-jensen *et al*. 2012; Kim *et al*. 2014; Dickinson *et al*. 2015; Paix *et al*. 2015; Dickinson and Goldstein 2016; Nance and Frokjaer-jensen 2019; Ghanta and Mello 2020; Nonet 2020; El Mouridi *et al*. 2022), which combined with optical transparency and a rapid life cycle, make this organism a uniquely powerful system for *in vivo* biology.

Key advances in *C. elegans* genome manipulation methods have focused on increasing the efficiency of targeted insertions (Paix *et al*. 2015; Ghanta and Mello 2020; Nonet 2020; El Mouridi *et al*. 2022), streamlining the process of creating homologous repair templates (Dickinson *et al*. 2015; Ghanta and Mello 2020; Demott *et al*. 2021), and reducing the effort required to screen for desired modifications (Kim *et al*. 2014; Dickinson *et al*. 2015; Dickinson and Goldstein 2016; Nance and Frokjaer-jensen 2019; El Mouridi *et al*. 2022). However, these methods all require precise microinjections to deliver genetic material into the microscopic core of the gonadal syncytium, which is a difficult skill to learn and master. Despite steady progress towards increasing the ease and efficiency of genetic methods, there have been no comparable advances in the process of germline microinjection itself. Indeed, using current techniques it is possible for an individual to generate large numbers of homologous repair templates in parallel at a rate that exceeds the capacity to perform the injections. Accordingly, the physical process of microinjection is a major bottleneck for generating novel *C. elegans* strains and is also a key barrier to entry for labs learning to use genome engineering approaches.

A common rate-limiting factor for *C. elegans* germline microinjections is transferring and positioning the worms at various stages of the procedure. Worms are first moved from an NGM plate to a microscope slide for injection, where they are immobilized by adhering to a dried agarose pad in a thin layer of halocarbon oil. For the most efficient injections, worms must be oriented uniformly such that both arms of the gonad are easily accessible to the microinjection needle without damaging the germline or other essential structures. Following injection, worms are released from the agarose pad using a droplet of aqueous buffer and transferred to an NGM plate for recovery. Worms are typically handled throughout this process using a standard platinum wire worm pick (Kadandale *et al*. 2009; Rieckher and Tavernarakis 2017; Ghanta *et al*. 2021), although an eyelash pick is used by some labs. Throughout this process, great care must be taken to avoid damaging the worms, while still moving and positioning them fast enough to avoid fatal desiccation. Unfortunately, standard methods limit users to moving small numbers of worms at a time, and the pick must be used precisely with skill and care to avoid injuring the worms.

Here, we report that a small paintbrush is a gentle and effective method to rapidly move large numbers of worms throughout the microinjection process. In paired trials, the paintbrush method significantly increased both injection speed and post-injection survival compared to standard methods, which on average almost tripled the rate of injections. We also found that the paintbrush method improved the ability of novices to perform key steps in the microinjection process. These results indicate that this method will be beneficial both for experienced personnel seeking to increase the speed of generating strains and new personnel learning to inject. Importantly, the paintbrush method is inherently compatible with all *C. elegans* genetic methods that rely on microinjections and can be implemented quickly and with minimal cost and effort. By lowering technical barriers, we expect the relative ease of our paintbrush method will accelerate the pace of generating new *C. elegans* lines and make genome manipulation approaches more widely accessible.

## Results and discussion

Microinjecting *C. elegans* usually involves three steps that require manually handling the worms under a stereomicroscope: (1) moving young adult hermaphrodites from an NGM plate into halocarbon oil; (2) mounting them onto a dried agarose pad where they are immobilized for injection; and (3) releasing the worms from the pad using M9 buffer and transferring them to a recovery plate after injection. The second two steps are critically time-sensitive, as the worms can only survive for several minutes on the dried agarose pad. Traditionally, animals are moved and positioned throughout this process using a metal worm pick (Kadandale *et al*. 2009; Rieckher and Tavernarakis 2017; Ghanta *et al*. 2021), although some labs use an eyelash pick. In addition to being difficult and often frustrating to learn, these methods limit the speed of injections because it can be challenging to precisely position multiple worms at a time.

In an effort to improve the ease and speed of *C. elegans* microinjections, we developed a simple and inexpensive strategy to move worms using small paintbrushes (10/0 – 3/0 sized) throughout the injection process (Fig. 1A, B; Videos S1 – S3). In our initial experiences, handling worms with a paintbrush rather than a pick qualitatively increased the ease and speed of worm handling at all steps and improved overall efficiency of the microinjection process. Experienced investigators who were originally trained to inject using traditional methods universally self-reported higher efficiency and improved worm survival with the paintbrush. In the first step of the injection process, we found that a small paintbrush dipped in halocarbon oil can rapidly pick up over 20 worms at a time from an NGM plate and easily release them into a drop of halocarbon oil on the injection cover slip (Video S1). In the next step, the paintbrush also helped with the ease and speed of transferring freely moving worms from halocarbon oil onto the agarose pad and immobilizing them in the ideal orientation for germline microinjections (Video S2). With the paintbrush, it was possible to routinely and precisely position 15 or more animals in a parallel line in less than one minute. Simply swiping the paintbrush in one direction oriented the worms in a rough line, and they could be furthered maneuvered individually with minimal risk of damage (Fig. 1A; Video S2). Use of a paintbrush for this step also facilitated removal of residual bacteria that can prevent the worm from adhering to the agarose pad or clog the microinjection needle. In the final injection step, worms are released from the agarose pad using a drop of aqueous buffer and transferred to a second NGM plate. In this step, the paintbrush made it possible to routinely transfer large numbers of worms (up to 10) at a time (Fig. 1B; Video S3).

**Figure 1.**
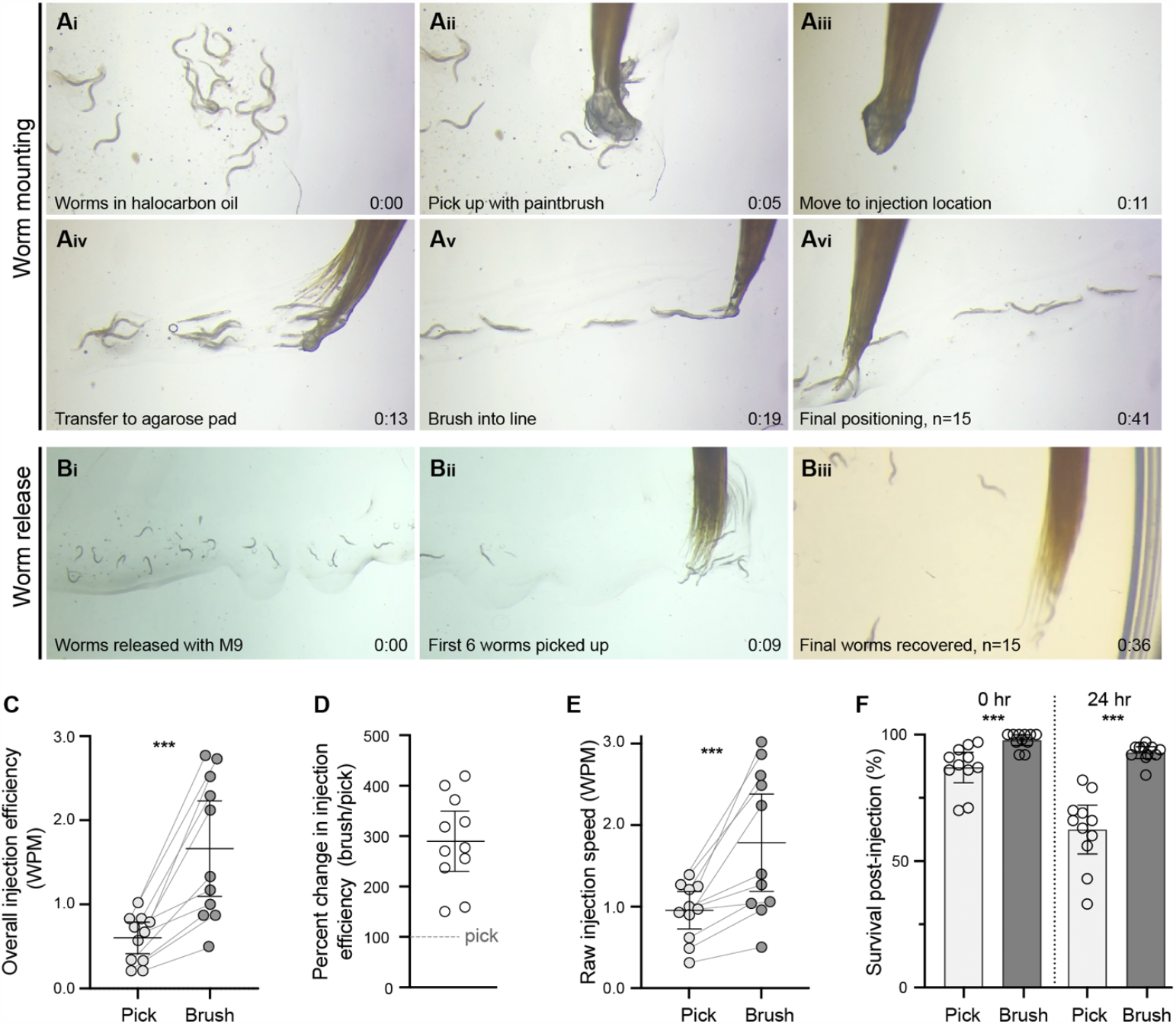
Using a paintbrush for worm handling dramatically improves *C. elegans* germline microinjection efficiency. (**A, B**) Representative images of key steps in the worm microinjection process as performed with a fine paintbrush. (**A**) Still images from Video S2 showing worms being moved from halocarbon oil and aligned on a dried agarose pad for microinjection within 41 seconds. (**B**) Still images from Video S3 showing 15 worms being safely recovered from the agarose pad to an NGM plate within 36 seconds. (**C** – **F**) Quantitative analyses of paired microinjection trials performed by experienced investigators using the paintbrush and traditional pick methods. The paintbrush method significantly increased overall injection efficiency compared to the traditional pick (**C**) (p=0.0010) resulting in an average 290% increase in microinjection throughput (**D**). This increase in overall efficiency can be attributed to both significantly increased speed (**E**) (p=0.0010) and survival after injection (**F**) (p*=*0.0010 at both time points). Error bars denote the mean and 95% confidence intervals. See Table S1 for raw data.

To confirm these improvements in injection efficiency quantitatively and investigate their underlying causes, we conducted paired time-trial experiments where experienced individuals used both a paintbrush and a pick during the same round of injections. For these experiments, experienced injectors attempted to inject as many worms as possible over a ∼30 minute period with either a metal pick or a paintbrush, followed by 30 minutes of injections using the other method, with the same needle if possible (Fig. 1C – F; Supplemental table 1). For each trial, we quantified the time elapsed, the total number of worms injected and recovered, the number of worms that survived the initial injections, and the number of surviving worms 24 hours after injection. We calculated overall injection efficiency for each trial in terms of worms per minute (WPM), which we defined as the number of surviving worms 24 hours after injection divided by the time spent injecting. Compared to the traditional method, the paintbrush method yielded robust and significant increases in overall injection efficiency (Fig. 1C, D; p=0.0010, Wilcoxon matched pairs signed rank test). Within paired trials, we found the efficiency of the paintbrush method was on average almost triple that of the traditional method (mean = 290%; Fig. 1D). Importantly, although we observed considerable variability in injection rates due to differences in individual proficiency and other factors, all paired trials showed within-pair increases in overall injection efficiency using the paintbrush compared to a pick (Fig. 1C, D). To identify the underlying cause(s) of these improvements in injection throughput, we compared raw injection speed (worms injected/time regardless of survival) and worm survival between the pick and brush methods. These comparisons showed that improvements in both categories contributed to the paintbrush method’s overall advantage in injection efficiency (Fig. 1E, F). The brush significantly increased raw injection speed (Fig. 1E; p=0.0010, Wilcoxon matched pairs signed rank test), and survival immediately after injection and 24 hours later (Fig. 1F; p=0.0010 at both time points, Wilcoxon matched pairs signed rank test). The difference in survival was most pronounced at 24 hours after injection when mean survival was 62% for the traditional method compared to 93% for the paintbrush method (Fig. 1F). Survival at this time point may be particularly relevant given that desired insertions using a common method for fluorescent protein knock-ins are most likely to occur more than 24 hours after injection (Ghanta and Mello 2020).

Because learning to microinject *C. elegans* using traditional methods can be extremely challenging for new personnel, we reasoned that the paintbrush method might also facilitate training novice researchers. As a proxy, we tested the extent to which using a paintbrush helped individuals who had not previously injected *C. elegans* to perform the key worm transfer steps require for germline microinjections. To do so, we tested the abilities of individuals without prior experience injecting *C. elegans* to move ten worms from an NGM plate to halocarbon oil, mount them on an agarose pad in a rough line, and recover them to a second NGM plate using both a traditional metal worm pick and a paintbrush. We recorded the time to complete this process along with the number of worms successful mounted and the number surviving 24 hours after recovery (Supplemental table 2). Similar to experienced injectors, the paintbrush method dramatically increased overall efficiency for novice investigators in completing these tasks (Fig. 2A, B) through a combination of increased speed (Fig. 2C) and higher survival rates (Fig. 2D).

**Figure 2.**
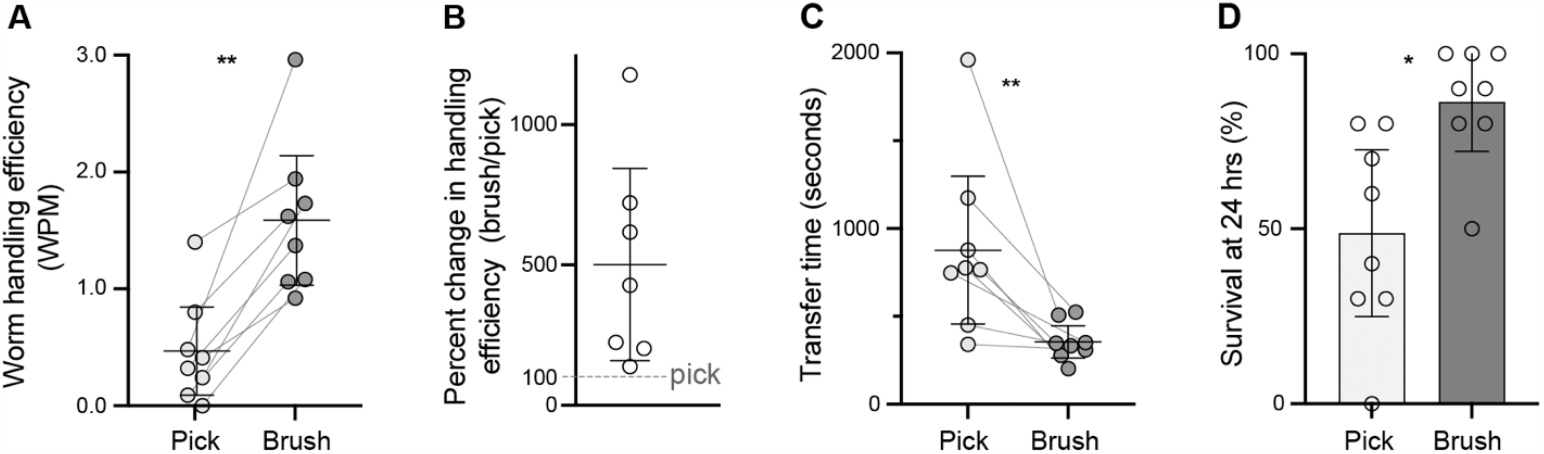
The paintbrush method facilitates worm handling by novice injectors. (**A** - **D**) Quantitative analyses of paired worm handling trials performed by individuals who had not previously attempted *C. elegans* germline microinjections. Compared to the traditional pick, the paintbrush method significantly increased worm handling efficiency (**A**) (p=0.0078) resulting in an average 501% improvement in efficiency (**B**). Similar to experienced injectors, novice injectors benefited from both faster worm handling (**C**) (p=0.0078) and increased survival (**D**) (p=0.0156) using the paintbrush method. Error bars denote the mean and 95% confidence intervals. See Table S2 for raw data.

## Conclusions

In comparison to traditional methods, our paintbrush method for worm handling dramatically increased germline microinjection efficiency by roughly tripling the number of worms that could be successfully injected in a given time. Individual injectors uniformly reported that the paintbrush method was faster and easier, and quantitative analyses confirmed that it significantly increased injection throughput through a combination of effects on microinjection speed and post-injection survival. We anticipate this simple and inexpensive method will accelerate the process of generating novel *C. elegans* strains using a variety of genome engineering approaches and help to make genome manipulation methods more widely accessible.

## Materials and Methods

### *C. elegans* handling

Briefly, our paintbrush method used 10/0 – 3/0 sized paintbrushes for all aspects of worm handling during the microinjection process. Separate paintbrushes were used for the worm mounting and recovery steps to ensure that aqueous M9 buffer was not present on the brush used for mounting worms on the dried agarose pad. We typically sterilize brushes in 70% or 100% EtOH between uses and wash them with soap and water between microinjection sessions. A detailed protocol for worm handling using a paintbrush for microinjection is provided in Supplemental Note 1, and at doi.org/10.17504/protocols.io.5qpvor1r9v4o/v1. Representative videos of key steps are presented in Videos S1 – S3. The worm handling process using a pick was similar to previously described methods (Ghanta *et al*. 2021) but varied slightly between experienced investigators based on their personal preferences.

### Microinjection method time trials

For the paired paintbrush and pick microinjection trials, we first moved an excess number of young adult worms onto an NGM plate with no bacteria using a flame-sterilized metal pick (this time was not included in the quantifications). Each time trial started with moving worms to inject from the NGM plate to a drop of halocarbon oil at one end of a 24 mm x 50 mm glass cover slip. Worms were then aligned for microinjections on a dried 2% agarose pad on the center of the same cover slip. Each investigator chose the number of worms to mount based on their prior experiences. After microinjecting both gonad arms for each animal, worms were immediately removed from the dried agarose pad using a drop of M9 buffer placed over the worms and transferred to an NGM plate with *E. coli* OP50 bacteria for initial recovery. After each paired trial, live worms were carefully moved from the recovery plates to new NGM/OP50 plates at a density of three worms per plate, and dead worms were counted. The number of dead worms was counted again 24 hours after microinjection to measure mortality occurring after the initial injection process. All worm manipulation steps were performed under a stereomicroscope (Nikon SMZ 1500 or Leica M80).

For the novice injector trials, approximately 50 worms were first moved to an empty NGM plate by experienced investigators. Following brief instruction on how to use the paintbrush and pick properly, novice injectors were then timed on completing the manual worm handling tasks required for germline microinjections. For each trial, we determined how long it took to move 10 worms from the NGM plate to a drop of halocarbon oil on a glass cover slip, how long it took to move worms from the halocarbon oil to a dried agarose pad, and how long it took to recover worms to a second NGM plate using both methods. Worm survival was assessed at 24 hours.

### Data analyses and figure preparation

Statistical analyses were performed using Graphpad Prism 9.5.1. All paired trial data were analyzed using two-tailed Wilcoxon matched pairs signed rank tests. Graphs were generated in Prism, and final figures were prepared using Adobe Illustrator 24.1. Videos were acquired using an Apple iPhone 6S with a GOSKY Smartphone Universal Adapter Mount (Amazon.com) and a 10x eyepiece on the second head Wild M8 stereo teaching microscope.

### Microinjections

*C. elegans* germline microinjections were performed on a Nikon Ts2R inverted microscope with floating stage, DIC optics, and 10x and 40x air objectives. We used a Narishige 3-axis oil-hydraulic micromanipulator (MMO-203) mounted on a Narishige coarse micromanipulator (MNM-4) attached to the microscope illumination column for needle positioning and a World Precision Instruments PV820 pressure injector.

### Key materials

Round-tipped paintbrushes sized 10/0 and 3/0. We typically use Craft Smart brand brushes purchased from Michaels, but any similarly sized brush with soft synthetic bristles can be used.

## Acknowledgments

We thank Alan Bergland, Amanda Gibson, Sarah Kucenas, Eyleen O’Rourke, and Sarah Siegrist for their support. We thank Jay Hirsh for use of the Wild teaching stereomicroscope for acquiring videos. We would like to thank Arun Dutta for helpful feedback on the manuscript draft. We also acknowledge the many groups who have published advancements in *C. elegans* genome engineering and transgenesis methods not cited here for reasons of space. Some strains were provided by the CGC, which is funded by NIH Office of Research Infrastructure Programs P40 OD010440.

## Author contributions

TVG: conceptualization, methodology, formal analysis, investigation, writing – original draft, visualization, project administration; AMP: formal analysis, investigation, resources, writing –review and editing, visualization, supervision, funding acquisition; MF, LL, TNM-K, DQM, DCM, ABA, KRB, LTB, AG, JAJ, KEL, CGM, RJP: investigation.

## Supplemental Videos

**Video S1. First-person perspective of transferring *C. elegans* from an NGM plate to halocarbon oil**

https://doi.org/10.6084/m9.figshare.22299274

Video depicts two representative examples of transferring *C. elegans* from an NGN to a drop of halocarbon oil at the edge of the injection cover slip. The first example starts at 0:05. Approximately 23 worms are picked up by 0:10. All worms are safely deposited in halocarbon oil by 0:17 with three swipes of the paintbrush. The second example starts at 0:22. Approximately 18 worms are picked up by 0:30 and safely deposited in two swipes in the halocarbon oil by 0:35. At 0:37, there is a final swipe that spreads the bristles to check if worms are still left in the bristles. Video corresponds to steps 5 and 6 in Supplemental Note 1.

**Video S2. First-person perspective of mounting *C. elegans* on dried agarose pad for microinjection**

https://doi.org/10.6084/m9.figshare.22320688

Video depicts two representative examples of moving *C. elegans* from the halocarbon oil worm reservoir to the agarose pad and orienting them for microinjection. The first example starts at 0:04. Swirling of the worms to remove excess oil and move the worms closer can be seen between 0:05-0:11. Approximately 17 worms are picked up in two swipes by 0:13. Mounting begins at 0:15, and the final worms are positioned by 0:52. Swiping away bacteria can be seen at 0:51. The second example begins at 0:56. Approximately 19 worms are swirled and picked up by 1:00, and are in final positions by 1:34 following repositioning. An individual worm being repositioned to better expose the gonad for microinjection can be seen at 1:30. Video corresponds to steps 7 and 8 in Supplemental Note 1.

**Video S3. First-person perspective of recovering *C. elegans* from agarose pad to an NGM plate**

https://doi.org/10.6084/m9.figshare.22320694

Video depicts two representative examples of recovering *C. elegans* from aqueous buffer on the injection cover slip and moving them to an NGM plate. Videos start with worms already released from the agarose pad with several drops of M9 buffer applied with a 20ul pipette just before the start of the video. The first example begins at 0:04. Following several swipes towards the bottom of the agarose pad to push the worms together, 6 worms are seen being picked up at once by 0:13. They are safely deposited onto an NGM recovery plate by 0:19. The remaining 9 worms are picked up and deposited in two rounds by 0:42. The second example begins at 0:48. 7 worms are initially picked up by 0:54 and deposited by 0:59. Approximately 10 more worms are picked up and deposited in two rounds by 1:27. A good example of swiping to ensure all worms are deposited and no worms remain stuck in the bristles occurs around 1:20-1:27. Video corresponds to steps 11 and 12 in Supplemental Note 1.

## Supplemental Tables

**Table S1.** Raw data for paired microinjection trials by experienced injectors.

| Trial | Time elapsed (min:sec) |  | Total worms injected |  | Overall injection efficiency (WPM)* |  | Change in efficiency** | Raw injection speed (WPM) |  | Initial survival |  | Survival at 24 hours |  |
| --- | --- | --- | --- | --- | --- | --- | --- | --- | --- | --- | --- | --- | --- |
|  | Pick | Brush | Pick | Brush | Pick | Brush |  | Pick | Brush | Pick | Brush | Pick | Brush |
| 1 | 28:46 | 33:11 | 14 | 32 | 0.21 | 0.87 | 419% | 0.49 | 0.96 | 86% | 97% | 43% | 91% |
| 2 | 30:00 | 30:00 | 30 | 42 | 0.33 | 1.33 | 400% | 1.00 | 1.40 | 87% | 100% | 33% | 95% |
| 3 | 30:00 | 30:00 | 28 | 38 | 0.73 | 1.17 | 159% | 0.93 | 1.27 | 96% | 100% | 79% | 92% |
| 4 | 30:00 | 32:00 | 29 | 34 | 0.67 | 1.00 | 150% | 0.97 | 1.06 | 97% | 100% | 69% | 94% |
| 5 | 36:05 | 27:15 | 34 | 109 | 1.03 | 2.77 | 270% | 1.25 | 3.02 | 91% | 92% | 82% | 92% |
| 6 | 30:10 | 30:32 | 42 | 76 | 0.83 | 2.29 | 277% | 1.39 | 2.49 | 71% | 92% | 60% | 92% |
| 7 | 33:21 | 32:35 | 30 | 73 | 0.57 | 2.12 | 372% | 0.90 | 2.24 | 70% | 97% | 63% | 95% |
| 8 | 25:15 | 21:56 | 32 | 63 | 0.83 | 2.73 | 328% | 1.27 | 2.87 | 88% | 98% | 66% | 95% |
| 9 | 23:56 | 22:12 | 29 | 58 | 0.79 | 2.52 | 318% | 1.21 | 2.61 | 86% | 100% | 66% | 97% |
| 10 | 29:14 | 29:48 | 18 | 31 | 0.34 | 0.87 | 255% | 0.62 | 1.04 | 94% | 100% | 56% | 84% |
| 11 | 34:00 | 39:00 | 11 | 20 | 0.21 | 0.49 | 237% | 0.32 | 0.51 | 91% | 100% | 70% | 95% |
| Mean |  |  | 27.00 | 52.36 | 0.59 | 1.65 | 290% | 0.94 | 1.77 | 87% | 98% | 62% | 93% |

Abbreviations: WPM, Worms Per Minute
\* Overall injection efficiency was calculated as the number of worms surviving at 24 hours divided by the time elapsed for worm handling and microinjecting.
\*\* Change in efficiency was calculated based on the overall injection efficiency for the brush/pick methods

**Table S2.** Raw data for paired worm handling trials by novice injectors.

| Trial | Time elapsed (seconds) |  | % Survival at 24 hrs |  | Worm handling efficiency (WPM)* |  | Change in efficiency |
| --- | --- | --- | --- | --- | --- | --- | --- |
|  | Brush | Pick | Brush | Pick | Brush | Pick |  |
| 1 | 277 | 775 | 50% | 0% | 1.08 | 0.00 | NA |
| 2 | 509 | 1961 | 90% | 30% | 1.06 | 0.09 | 1178% |
| 3 | 309 | 341 | 100% | 80% | 1.94 | 1.41 | 138% |
| 4 | 332 | 450 | 90% | 60% | 1.62 | 0.80 | 203% |
| 5 | 346 | 766 | 100% | 30% | 1.73 | 0.24 | 721% |
| 6 | 524 | 1174 | 80% | 80% | 0.92 | 0.41 | 224% |
| 7 | 202 | 878 | 100% | 70% | 2.96 | 0.48 | 617% |
| 8 | 350 | 748 | 80% | 40% | 1.37 | 0.32 | 428% |
| Mean | 356 | 887 | 86% | 49% | 1.59 | 0.47 | 501% |

Abbreviations: WPM, Worms Per Minute
\* Worm handling efficiency was calculated as the number of worms surviving at 24 hours divided by the time elapsed for worm handling.

## Supplemental Methods

### Supplemental Note 1

#### Protocol for handling *C. elegans* using a paintbrush for germline microinjections

##### Overview

This protocol describes a method for using a paintbrush to move *C. elegans* throughout the microinjection process. This method accelerates the microinjection process compared to traditional methods and improves the likelihood of worm survival. This protocol only describes the stages of the microinjection process that pertain to worm handling under a stereomicroscope. Please consult Videos S1 – S3 for visual depictions of these methods. For a description of other aspects of the microinjection process including the injections themselves, please see (Kadandale *et al*. 2009; Rieckher and Tavernarakis 2017; Ghanta *et al*. 2021).

##### Required materials

- Round tipped paintbrushes sized 10/0 and 3/0. We typically use Craft Smart brand brushes purchased from Michaels.

> *Note*: Different paintbrush brands and sizes can be used depending on personal preferences and availability. We recommend trying sizes between 30/0 – 1/0. In our experience, a 10/0 size brush is ideal for mounting worms, and a 3/0 brush is ideal for recovering them after injections. Wider brushes may be successfully used to recover worms after injections but are challenging to use for precision manipulations. For all steps, the bristles should be soft and fine to avoid injuring the worms.
- Cover slips with dried agarose pad for microinjections:
  - 24 mm x 50 mm #1.5 cover glass, Ephredia Richard-Allan Scientific #152450
  - Agarose, (2% in water spotted in middle of cover glass and dried overnight at room temperature)
- Halocarbon oil 700, Sigma Aldrich #H889850ML
- M9 buffer

##### Protocol

###### Move worms to inject from NGM plate to halocarbon oil

See Video S1, https://doi.org/10.6084/m9.figshare.22299274

1. Prepare well-fed, young adult hermaphrodite animals for injections by removing excess bacteria. We generally accomplish this by moving an excess number of worms to an NGM plate with no bacteria and allowing them to crawl for several minutes. Residual bacteria can be removed during later steps if necessary. Alternatively, worms can be picked into a dish of buffer to remove bacteria prior to mounting.
2. In the meantime, sterilize the mounting paintbrush (we use size 10/0) by dipping it in 70% ethanol. Allow to dry completely (about one minute).

> *Note:* This only needs to be done once at the beginning of the procedure, or before injecting new strains or injection mixes, but can be repeated throughout if contamination is observed. Using 100% ethanol will accelerate the brush drying if desired. We have found that contamination levels on post-injection plates are similar regardless of whether the injector utilized a paintbrush or flame-sterilized metal pick for worm handling.
3. Sterilize a separate paintbrush (we use size 3/0) to use for recovering worms after injection at the same time so that it is ready when needed.
4. Place a drop of halocarbon oil 700 (about 50 µL) onto one end of a 24 × 50 mm glass coverslip with a dried 2% agarose pad in the middle. This oil drop is where worms are pre-staged prior to injections to provide an easily accessible supply of worms that can be rapidly moved to the agarose pad for several injection rounds without needing to return to the NGM plate.
5. To move worms from the clean NGM plate (step 1) to the oil droplet, first dip the very tip of the mounting paintbrush into halocarbon oil. Then pick up one, or many, worms at a time by swiping over them or tapping on top of them. One can easily transfer 10-20 animals at a time.

> *Note:* Use caution not to embed worms deeply into the paintbrush bristles when picking them up. One should ideally use only the tip of the brush.
6. Dip the brush carrying worms back into the halocarbon oil drop at the edge of the glass cover slip (from step 4). Move the brush in the oil to release the worms. Worms should be freely moving in the oil drop.

> *Note:* If a worm is stuck, one can gently brush the bristles while rotating the brush directly on the cover slip to release the worm without causing damage (see Video S2). Any remaining bacteria can also be removed at this point by swirling the brush.

###### Orient and immobilize worms on the agarose pad in a line for injections

See Video S2, https://doi.org/10.6084/m9.figshare.22320688

7. Using the same mounting paintbrush previously dipped in halocarbon oil, pick up the desired number of worms to inject with the tip of the paintbrush.

> *Note*: One can grab 1-20 worms at this point depending on preference and experience level. It is ideal to first rotate and twist the brush on an empty part of the coverslip to remove any excess oil stuck in the bristles prior to this step as too much oil can complicate immobilizing the worms. When grabbing worms in the halocarbon oil, you can reduce excess oil and grab more worms by swirling the brush around several worms to push them together.
8. Position the paintbrush with worms near the top of the agarose pad where you wish to inject. Brush downwards to start positioning the worms in a line. Use the brush to move worms to the desired locations and orient them for injection.

> *Note*: Using the paintbrush greatly improves the ability to reposition worms on the agarose pad without damage compared to a metal or eyelash pick. If needed, one can brush over a worm several times to ensure it sticks to the agarose pad in the ideal orientation for germline microinjection. If a worm is not sticking well, one can move it to another section of the agarose pad or wait a few seconds before brushing it against the agarose pad again. It is important to ensure that all worms are completely covered in halocarbon oil after they are immobilized to limit desiccation and to avoid air/oil interfaces near the worm that interfere with viewing the needle tip and germline simultaneously on the compound microscope used for injections. If needed, one can add a small amount of halocarbon oil to the tip of the brush and dab oil on any worms that require more without hurting them.
9. Perform germline microinjections using standard procedures.

###### Release worms from agarose pad and transfer them to NGM plate

See Video S3, https://doi.org/10.6084/m9.figshare.22320694

10. After injecting, add an aqueous buffer, such as M9, in 2-3 µL drops over the worms to release them from the agarose pad. The worms will generally float off the pad on their own. Using the minimum amount of M9 required will make it easier to transfer multiple worms at a time.
11. Using the recovery paintbrush (we use size 3/0), swipe the worms to capture them with the tip of the bristles. Make sure to *use separate paintbrushes for this step and for mounting worms*. Any aqueous buffer on the mounting paintbrush can prevent the worms from sticking to the agarose pad.

> *Note*: To recover multiple worms at a time, guide several worms towards the edge of the M9 droplet before quickly swiping while rotating the paintbrush. With practice, one can grab at least ten or twenty worms at a time. It may be helpful to count the worms at this point to ensure all are recovered in the next step.
12. Transfer the worms to an NGM plate with *E. coli* OP50 bacteria for recovery by swiping the paintbrush onto a bacteria-free section of the plate.

> *Note*: All worms should be deposited on the NGM plate after a few brush strokes. If a worm is stuck, flip the brush, or twist it continuously, and rapidly move it on the plate until the worm is released.
13. When finished, clean the brushes by washing them gently with soap and water and rinsing with distilled water. Ensure all soap is washed out and reshape the bristles into a fine tip. Allow to air dry.

